# Natural slab photonic crystals and where to find them among the girdle bands of diatoms

**DOI:** 10.1101/2024.03.08.584047

**Authors:** Matt P. Ashworth, Daryl W. Lam, Martin Lopez-Garcia, Schonna R. Manning, Johannes W. Goessling

**Affiliations:** UTEX Culture Collection of Algae, University of Texas at Austin, Austin, TX USA; Department of Biological Sciences, University of Alabama, Tuscaloosa, AL USA; Instituto de Òptica, Consejo Superior de Investigaciones Cientìficas (IO-CSIC), Serrano 121, Madrid, Spain; Institute of Environment, Department of Biological Sciences, Florida International University, Miami, FL USA; Laboratory for Innovation and Sustainability of Marine Biological Resources (ECOMARE), Centre for Environmental and Marine Studies (CESAM), Department of Biology, University of Aveiro, Portugal

**Keywords:** Slab photonic crystals, Diatoms, Correlated evolution, Trait evolution, Functional morphology

## Abstract

Slab photonic crystals, nanomaterials characterized by periodic pores for manipulating light, have applications in advanced optical technologies. Remarkably, similar materials have been identified in the silica shell of diatoms, in particular the girdle bands. Despite the potential applications and significance for diatom biology, their prevalence remains uncertain due to limited observations across a few species. In this study of 393 SEM girdle band micrographs across major taxonomic groups, we identified slab photonic crystals using Fast Fourier Transform (FFT) analysis. A correlation analysis of these properties on a phylogenetic tree revealed their distribution across the diversity of species and taxonomic groups. Square and hexagonal lattice varieties are prevalent in earlier-diverging groups, and linked to phytoplanktonic lifestyles. More recently-diverged clades lack these structures entirely in their girdle bands. Numerical analysis indicates that square lattice types exhibit anticipated photonic properties (stopbands) in the visible spectrum, while hexagonal lattice types are primarily linked to the near to mid-infrared range. This suggests that girdle band slab photonic crystal morphologies 1) originate from quasi-periodic photonic structures, 2) are primarily found in evolutionarily older clades (Coscinodiscophyceae and Mediophyceae), 3) lost square lattice types through diversification in the Mediophyceae, and 4) are absent in more recent clades (Fragilariophyceae and Bacillariophyceae). The limited inter-species distribution of slab photonic crystals may offer experimental cues to study their biological functionality. While these data suggest that stopband functionalities are a derived frustule trait, the ultimate purpose of slab photonic crystals in nature remains a mystery.

## Introduction

Geometry is a common phenomenon in nature, serving to optimize various biological processes including energy distribution, nutrient absorption, and growth^1,2^. One captivating aspect of geometry is periodicity, which involves the repetition of patterns at regular intervals. Periodicity can be observed across different scales, from the macroscopic to the nanoscale. At the nanoscale, periodic structures can introduce characters such as photonic properties, which enable the manipulation of light. Recent experiments have confirmed the presence of advanced photonic properties within the glass shell (frustule) of a few diatom species, bearing striking resemblances to the modern concept of slab photonic crystals^3^.

Photonic crystals are periodic nanoscale structures with profound implications for some of the most advanced modern technologies, such as optical communications, high-performance lasers, efficient solar cells, ultra-sensitive sensors, quantum information processing, and more. Out of the wide range of photonic crystal morphologies, the slab photonic crystal represents the most useful one by enabling intricate light manipulation in up to three dimensions with relatively simple configurations^4^. The concepts facilitating the nanofabrication of slab photonic crystals were initially developed in the late 1980s, opening new avenues for the precise engineering of light at nanoscale. However, more recently it was discovered that structures reminiscent of slab photonic crystals also exist in nature, in the diatom shell, as proposed through numerical simulations^5^ and later confirmed by experimental observations^3^. Diatoms (Bacillariophyta), which diverged between 200-182 million years ago^6^, predate human advancements in light manipulation for technology by hundreds of millions of years. However, at present, there are few hypotheses that explain the presence of these sophisticated photonic materials in diatoms. Also, there is no clear understanding of how prevalent these structures are across the diversity of diatoms or if they could potentially exist as byproducts in only a handful of species without specific links to their biology and ecology.

The diatoms are a photosynthetic microeukaryote group characterized by their distinctive silica walls, called frustules. The frustule consists of two overlapping thecae, each composed of a valve and one or a series of overlapping girdle bands linking the valves to the other theca. Beyond these broad characters, the frustules vary in overall and in fine morphology across the diversity of diatoms (Supplemental Fig. S1). Frustules have long served as the basis for the taxonomic classification of diatoms. As the fine morphology of the frustule correlates to species identification in a group of organisms where species diversity is estimated to be in the tens of thousands or more^7^, there is significant variability in the distribution, density and morphology of the pores which perforate the frustule. Among various proposed applications of these natural nanomaterials^8,9,10^, the frustule, or at least parts of it, has also been considered as a photonic platform, where naturally occurring structural periodicity might drive the manipulation of light at nanometer scales.

Researchers have previously documented some optical characteristics of frustules cleared of their organic components, as phenomena like iridescence become visible in dark field light microscopy^11^. Under certain lighting conditions, elements can appear colored as a consequence of angular illumination despite the colorless nature of the frustules’ silica core material^12,13^. Such photonic properties can manifest through various mechanisms, including light diffraction, scattering, or through refraction and reflection over porous features. The foundation of these properties lies in the refractive index contrast between the photonic structure and the surrounding medium with a different refractive index. The number of repeated events over which such period exists builds the photonic response and controls its intensity. While a single line of periodic pores (see Supplemental Fig. S1C) can already possess some photonic properties, slab photonic crystals are characterized by periodic morphologies in 2 or 3 dimensions (Supplemental Fig. S1D). The photonic response of photonic crystals in the UV-VIS spectral range is due to light interference induced by features of a few hundred nanometers, similar to the wavelength of the visible light. The biosilica of the valve and girdle band silica slabs presents periodically arranged structural features like pores which are filled with a different material, facilitating refractive index contrast. While the valve portion of the frustule might be most frequently documented as a photonic nanostructure, their properties as photonic crystals have been debated due to a potentially more complex structure, which differs from the common slab photonic crystal morphology. The girdle bands of some taxa show better defined photonic crystal slab morphologies than the corresponding valves^3^. These girdle bands are likely to be the preferred unit for photonic applications, as their overall perforation tends to be more uniform than that of valves and they lack additional ornamentation, such as siliceous crosswalls, elevated pore fields or siliceous spines found in valves.

Girdle bands can exhibit intricate morphologies in some diatom taxa, featuring a porous lattice in three dimensions, which bears a striking resemblance to advanced photonic components such as the previously mentioned slab photonic crystals^14^. Remarkably, it remains the sole nanostructure with a comparable architecture known in nature. Different girdle pore lattice types exist, including hexagonal type and square types^3^, which may allow for versatile light manipulation and control. The spectral response is primarily linked to the photonic crystal morphology including pore-to-pore distance (period) and pore diameter, the latter of which defines the volume into which the contrast medium can be inserted^15^. Despite such advanced photonic properties and potentially natural preservation, currently there is no profound hypothesis explaining how the girdle slab photonic crystal system could contribute to the physiological functions of live diatoms.

While there is a growing interest in exploring the photonic properties of the diatom frustule, whether for the aforementioned applications or to comprehend its role within the organism, these investigations have understandably concentrated on a limited number of diatom genera. Yet, the frustule morphologies studied so far represent just a fraction of the diverse morphologies found among diatoms. Thus, hypotheses about the functional implications of the diatom frustule based on the presumed photonic properties of these tested strains, such as the attenuation of UV radiation^16,17^, or the manipulation of light to reduce the formation of oxygen radicals^18^ or to increase photosynthetic efficiency^19,20,21^, should really be limited to taxa with the specific tested morphologies, rather than broadly applied to all diatoms. Only a few studies have focused on girdle bands, finding that the tested species feature precisely defined uniform lattices of pores on their girdle bands, suggesting slab photonic crystal properties. In particular, the lattice of pores in the girdle bands of *Coscinodiscus granii* creates a so-called photonic stopband; that is, a spectral band for which light propagation within the nanostructure is prevented, in this case for light in the near infrared spectral range, when immersed in water. On the other hand, light propagation in the green spectral range is promoted *via* waveguiding effects when the structure is immersed in water or other low refractive index contrast media^3^.

If these slab photonic crystals provide some selective advantage, we would expect these properties to be common across diatoms, or at least trigger intense diversification in the clade where these properties evolved. Alternatively, if the selective advantage of photonic properties are limited, or restricted to a certain habitat, we would expect such lattices in only a few clades. To that end, here we have documented the ultrastructural characteristics of the girdle elements for hundreds of diatom species across the four broad taxonomic categories (Coscinodiscophyceae, Mediophyceae, Fragilariophyceae and Bacillariophyceae; Supplemental Fig. S2). Using a model developed to identify photonic crystals based on the periodicity and density of pores in frustule components^3^ we screened a dataset of SEM micrographs from 393 strains across the taxa to determine just how common photonic crystals in the girdle bands are among the diatoms. The evaluation of frustule morphology and photonic model parameters, alongside a molecular phylogeny of the examined diatoms, aimed to identify any patterns of correlation between photonic crystals in the girdle bands and particular clades or ecologies.

## Results

It is well established that certain lattice-like arrangements and densities of pores are required for slab photonic crystal properties^22^. For the girdle band to show slab photonic crystal properties, the silica frustule should present pores arranged in an ordered or quasi-ordered, 2- or 3-D lattice (see the Materials section for definition of quasi-order). These lattice structures are unevenly distributed across the diversity of diatom girdle bands (Fig. 1). Among the tested strains, taxa with potential photonic girdle morphologies are exclusively found in the Coscinodiscophyceae and Mediophyceae classes, entirely absent in both the Fragilariophyceae (“araphid pennates”) and Bacillariophyceae (“raphid pennates”) subclasses. These lattice structures appear in two broad categories: hexagonal and square lattices. While both are present in the Coscinodiscophyceae and Mediophyceae, the square lattice represents a larger proportion of the coscinodiscophytes while the mediophytes overwhelmingly exhibit hexagonal lattices.

**Figure 1.**
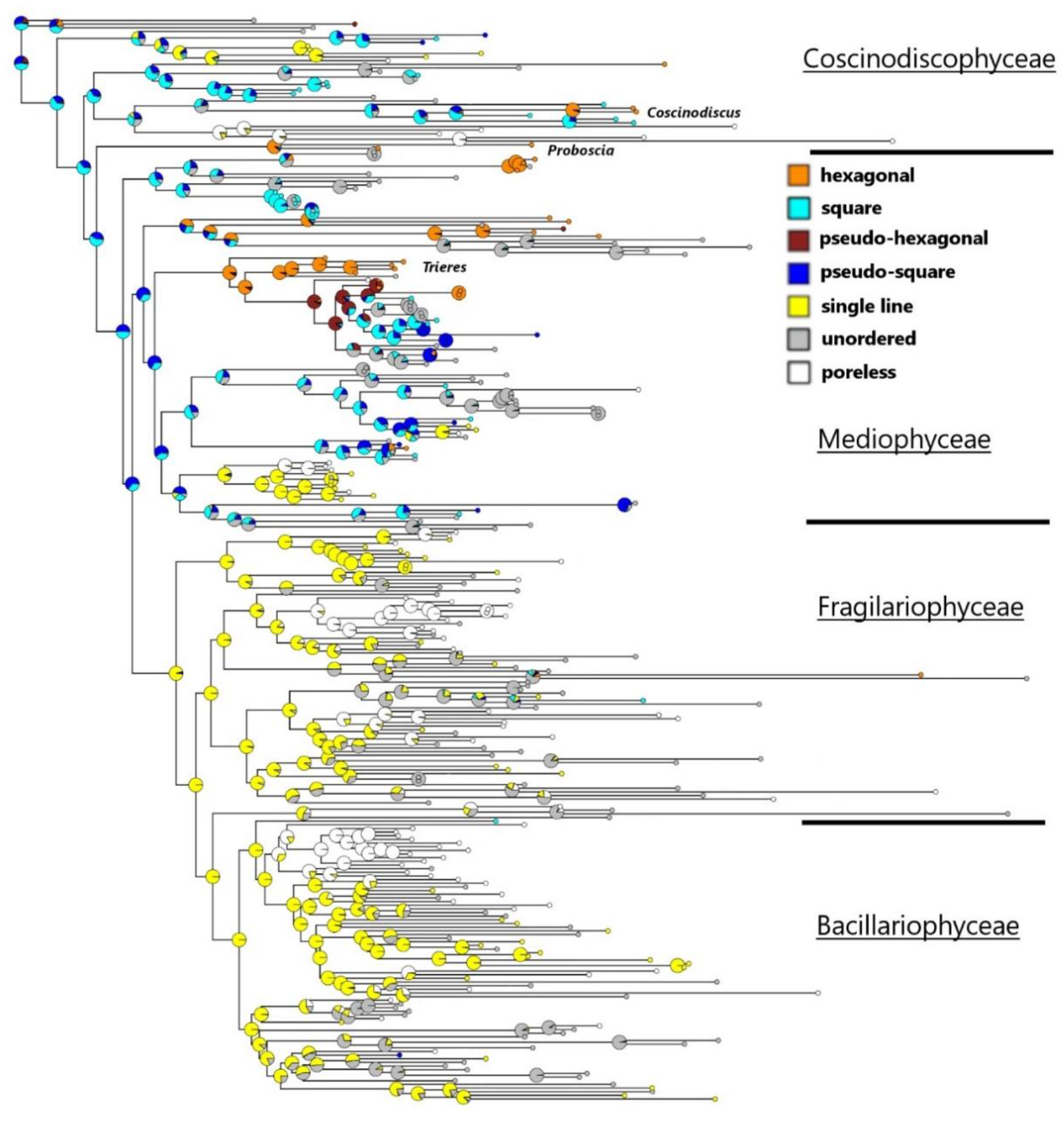
Pore morphology distribution in girdle bands across diatom phylogeny. Phylogenetic tree based on DNA sequence data from the diatoms evaluated for photonic girdle bands. The likelihood of the distributions of specific lattice patterns in the girdle band poration are indicated at the nodes and branch tips. These lattice patterning states are labeled per the key provided at the right of the tree figure. The limits of the major taxonomic groupings are also illustrated at the right of the tree. Genera relevant to Figure 2 are also included for reference.

There is a distinctive shift towards unordered perforation, single row or poreless girdles at the Bacillariophyceae (Fig. 1). Within the coscinodiscophycean clades, square lattices are present at higher relative abundance in the Rhizosoleniaceae (*Rhizosolenia* and *Guinardia* H.Peragallo) and Coscinodiscus clades. Each of these clades do feature taxa with hexagonal lattices as well (*Pseudosolenia calcar-avis* [Schultze] Sundström; *Coscinodiscus wailesii* Gran & Angst and *Coscinodiscus* sp. HK298, respectively). It should be noted that while the Rhizosoleniaceae features taxa with lattice structures on their girdle elements, no slab photonic crystal candidates were identified among the strains. The genera *Proboscia* and *Pseudosolenia* Sundström appear to be the sole genera among the coscinodiscophycean clades which exclusively feature hexagonal lattices. Among the mediophycean clades, two clades feature an abundance of lattice structures of both types. The aforementioned slab photonic crystal candidate *Trieres* is part of a larger clade of taxa in the Odontellaceae which contains additional genera bearing square (*Ralfsiella* [Ralfs] Sims, Williams & Ashworth) and hexagonal (*Pleurosira* [Meneghini] Trevisan) lattices. The genus *Biddulphia* Gray features both lattices, with *B. biddulphiana* (Smith) Boyer exhibiting a hexagonal lattice and multiple square lattices in the *B. alternans* (Bailey) Van Heurck/*B. reticulum* (Ehrenberg) Boyer clade. Hexagonal lattices are also found in *Trigonium* Cleve, *Biddulphiopsis* Stosch & Simonsen and *Lampriscus* Schmidt.

With regard to the cross-sectional ultrastructure of the girdle elements, all of the slab photonic crystal candidates have loculate frustules, those types that possess chambers or compartments (Fig.2). Porose frustules dominate the phylogeny, while loculate frustules are distributed unevenly, concentrated highest in the Coscinodiscophyceae and only a few genera in the Mediophyceae (*Neocalyptrella* Hernández-Becerril & Meave, *Trieres, Trigonium*) and Bacillariophycidae (*Brachysira* Kützing, *Parlibellus* Cox). As with the lattice structures, however, not all taxa with loculate frustules are slab photonic crystal candidates, including those within the Rhizosoleniaceae.

On a finer scale, girdle elements with potential photonic crystal properties are found in a very small fraction of the sampled diatom diversity (Figure 2). In fact, only three clades were exclusively composed of taxa exhibiting slab photonic crystal properties in their girdle elements: the *Coscinodiscus* and *Proboscia* Sundström clades (Coscinodiscophyceae) and the *Trieres* Ashworth & Theriot+*Triceratium spinosum* Bailey clade (Mediophyceae). There were a few other slab photonic crystal candidates in the Coscinodiscophyceae (*Hyalodiscus stelliger* Bailey) and Mediophyceae (*Triceratium robertsianum* Greville and *Thalassiosira pseudonana* Hasle & Heimdal), but these were outliers among the other taxa in their respective clades.

**Figure 2.**
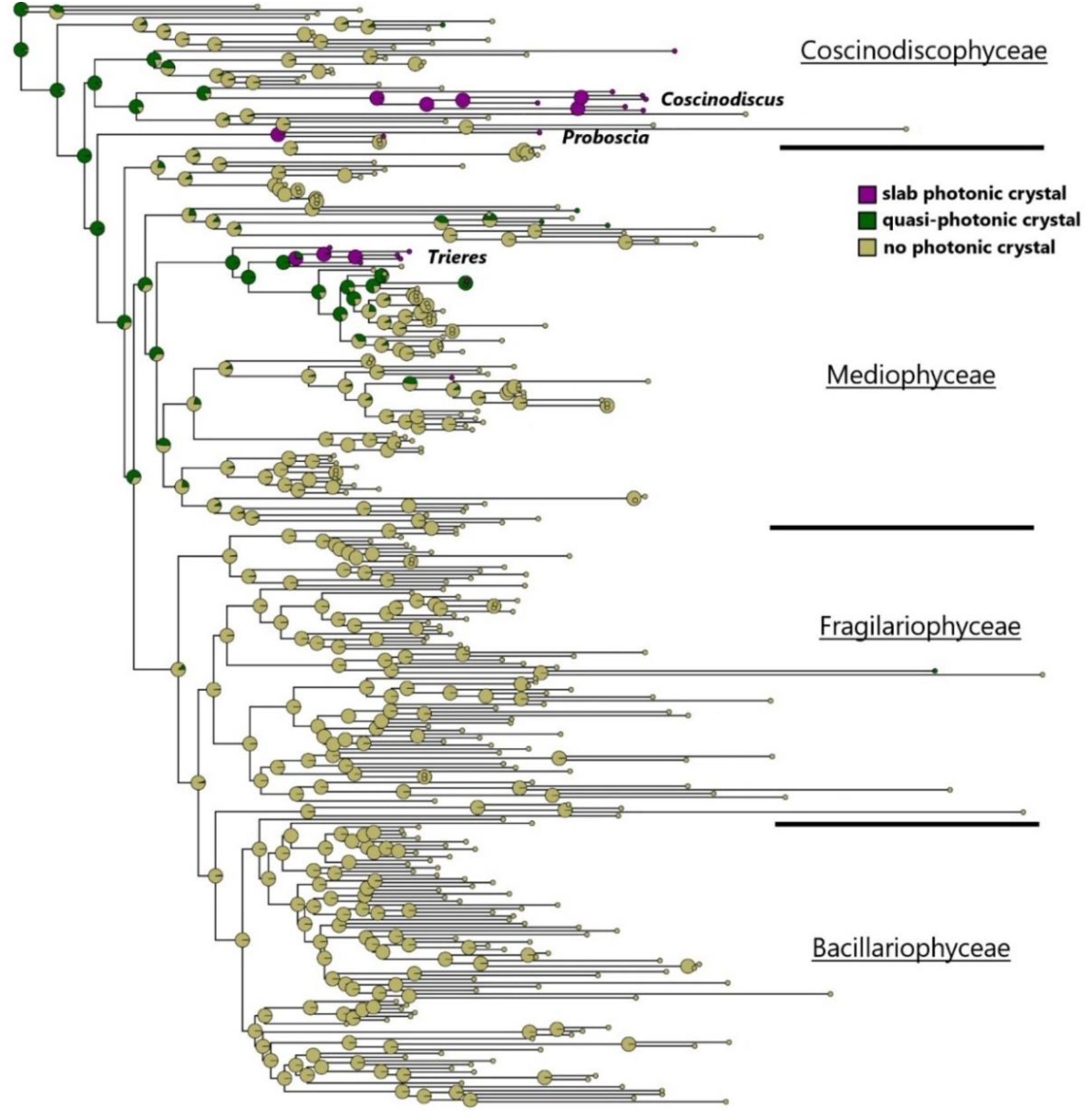
Phylogenetic tree based on DNA sequence data from the diatoms evaluated for photonic girdle bands. The likelihood of photonic properties in the girdle bands are indicated at the nodes and branch tips. The limits of the major taxonomic groupings are illustrated at the right of the tree. Only the genera in which all tested taxa show photonic girdle bands are labeled in the tree. A tree diagram which includes all taxon labels is provided as Supplemental Figure S3.

For taxa exhibiting slab photonic crystal-like lattice morphologies, analyzing the lattice parameter (period in x-direction) is crucial for inferring potential photonic properties and the spectral range using well-established models^3,15^. As depicted in Figure 3, species with square or quasi-square lattices generally exhibit lattice parameters ranging from 100 to 300 nanometers, pertinent scales for eliciting photonic properties in the visible to near-infrared spectral range (450–1000 nm), according to models developed by the authors^15^. Figure 3 further illustrates the anticipated central wavelengths for the photonic stopband relative to the lattice parameter defined above. The stopband defines the spectral range in which light is impeded from propagating within the girdle band in this case along the ΓM directions for both hexagonal and square lattices (z-direction in Fig. 4A)^3^. Notably, the central wavelength falls within the visible to near-infrared range for most species with a square lattice. Some species display larger values, resulting in significantly longer operating wavelengths for photonic properties. For instance, in the case of *Ralfsiella smithii* (HK331 in Fig. 3A) with a value as large as lattice parameter ≈ 1000 nm, this corresponds to a central wavelength in the infrared regime (λ ≈ 3200 nm). Interestingly, hexagonal-type lattices (Fig. 3B) exhibit a heterogeneous distribution of lattice parameters, ranging from lattice parameter ≈ 100 to ≈ 2000 nm with no apparent preferential range of values. This variability is reflected in a wide range of central operation wavelengths for the photonic properties, covering a slightly broader spectrum than square lattices, extending up to 4000 nm wavelengths (mid-infrared).

**Figure 3.**
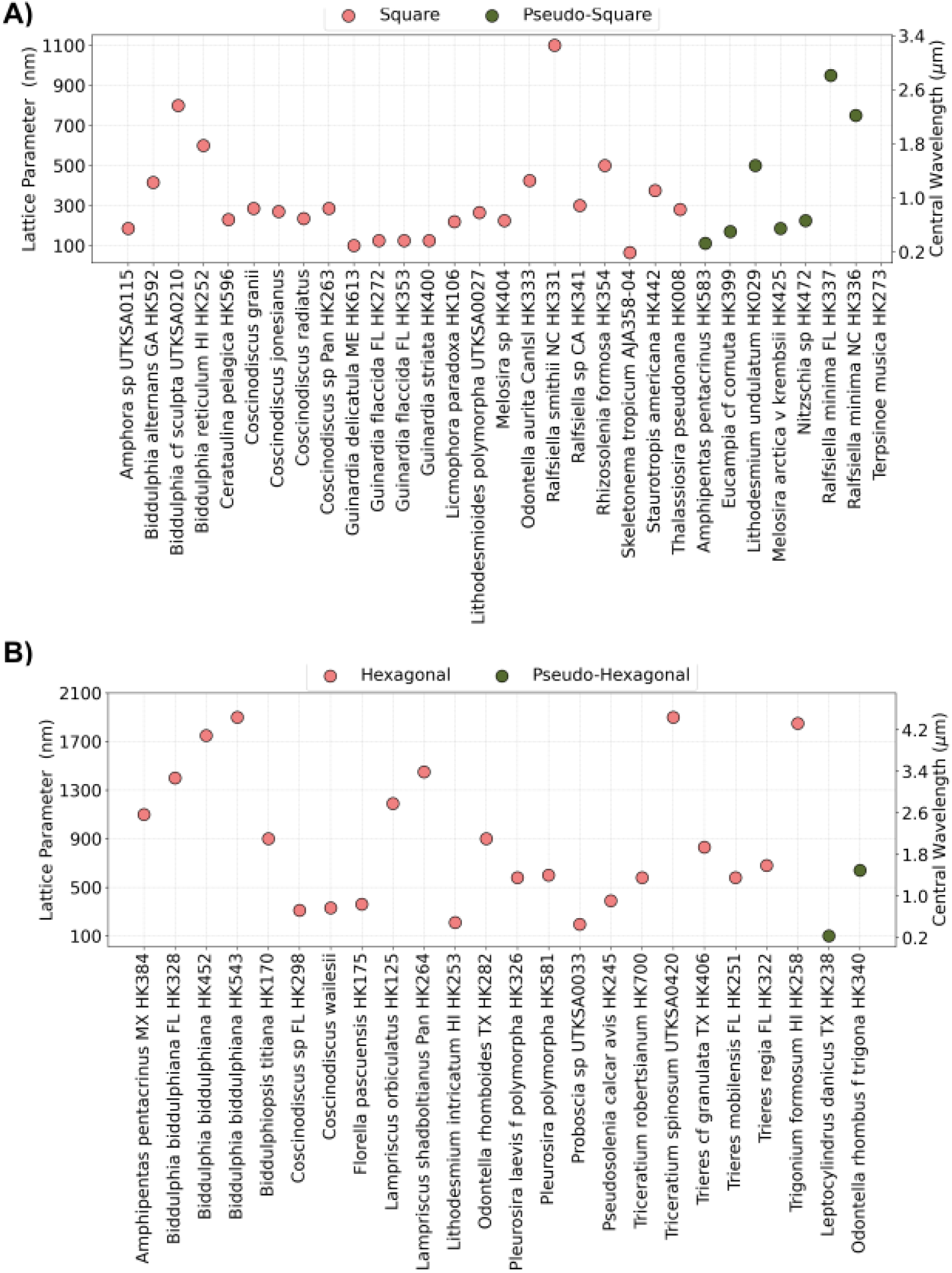
Expected Spectral Properties of Photonic Stopbands in Different Species. Distribution of lattice parameters (pore-to-pore distance) and central wavelengths of the photonic stopbands in girdle bands identified as photonic crystals for: A) Square and pseudo-square types. B) Hexagonal and pseudo-hexagonal types.

**Figure 4.**
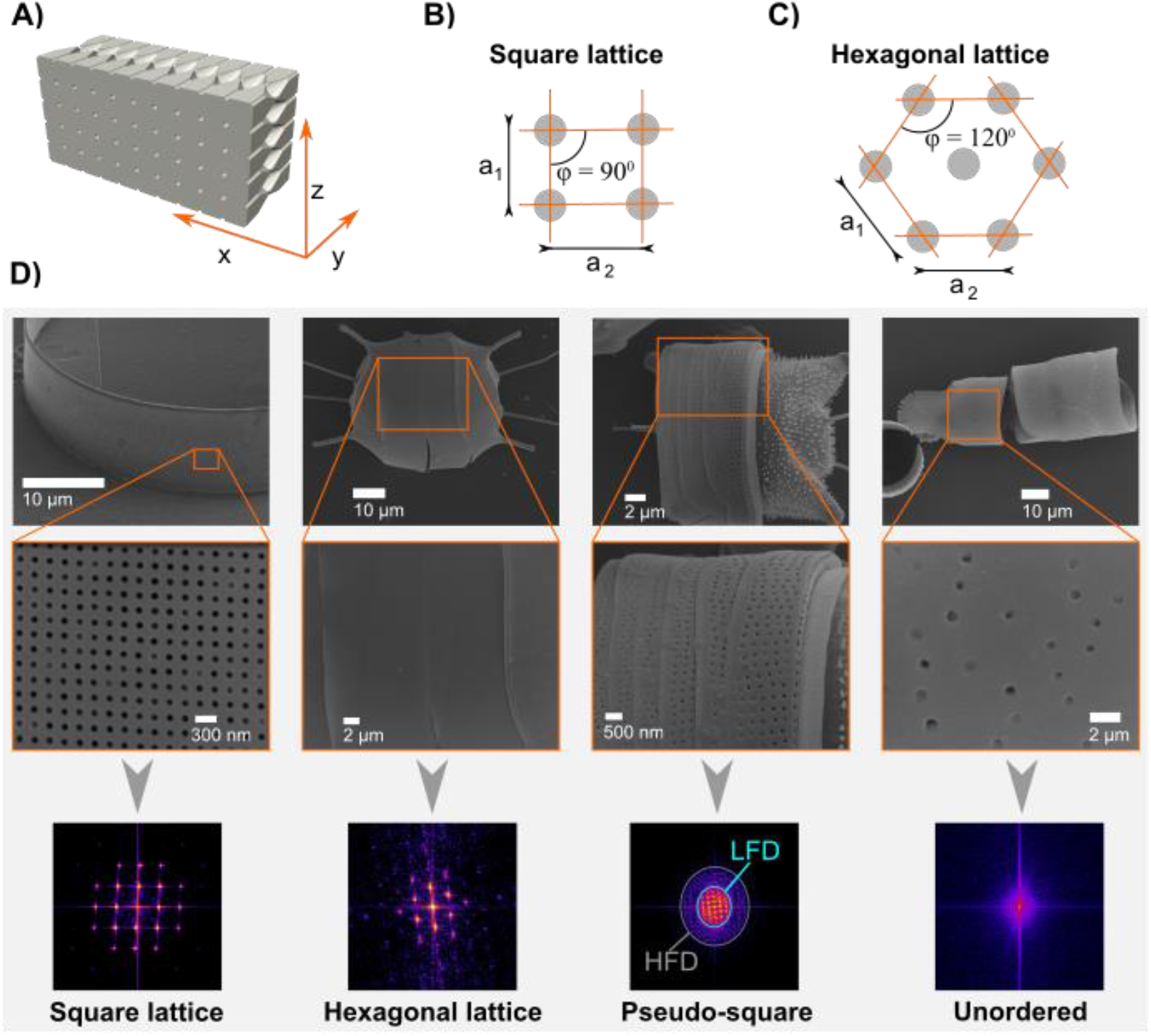
Determination of lattice symmetries in the girdle bands of diatoms. A) Illustration defining directions along the expected pore symmetry in diatom girdles, reminiscent of slab photonic crystals. Notably, the internal rhombic chamber system (indicated) was previously observed in species *C. granii* but is not considered in this analysis. B) Definition of square lattice symmetry, and definition of hexagonal lattice symmetry with two lattice vectors (a_1_ and a_2_) and the angle (φ) between them. D) Examples of Fast Fourier Transform (FFT) analysis performed on SEM micrographs. Symmetrical patterns in the low-frequency domains (LFD) indicate high structural symmetry, while high-frequency domains (HFD) suggest disorder. Species, from left to right: *Coscinodiscus sp*., *Trieres mobiliensis, Odontella sp*., *Orthoseira dendroteres*.

## Discussion

The extensive variety of diatoms and their diverse frustule shapes have posed challenges to our understanding of photonic properties and how they may function in nature. In this study, we focused on exploring the presence of slab photonic crystal properties within the girdle bands, one part of the diatom frustule, examining both their diversity and taxonomic distribution. The photonic crystal properties emerge from highly geometrical ornamentation on the nanoscale, which we estimated using FFT image analysis over SEM micrographs. Our investigation revealed that slab photonic crystal properties are primarily confined to three clades of phytoplanktonic diatoms belonging to the relatively older evolutionary taxonomic groups: the Coscinodiscophyceae and Mediophyceae. The more evolutionarily derived “pennate” clades, at least those tested in this study, did not exhibit slab photonic crystal morphologies within their girdle bands.

Our phylogenetic analysis provides insight into the emergence of highly ordered materials, exemplified by the slab photonic crystals, originating from quasi-ordered photonic structures positioned at the base of the phylogenetic tree (Fig. 1). In this context, we define quasi-photonic crystal morphologies, exhibiting either square or hexagonal characteristics, as those manifesting weak lattice symmetry identified through the presence of high-frequency domains in FFT analysis^23^. Noteworthy is the acknowledgment that these quasi-photonic structures may impart photonic properties, albeit with anticipated differences in the intensity and spectral confinement of these properties compared to slab photonic crystals. The influence of quasi-symmetry extends to potential asymmetry, resulting in the complete loss of pore features within the most derived evolutionary clades, specifically Fragilariophycecae and Bacilariophyceae. In summary, the trajectory of quasi-order takes two divergent paths: it either evolved into 1) highly ordered slab photonic crystals, or 2) disordered lattice symmetries, which thereafter evolved into pore-less structures in the most recent groups of diatoms. However, once a slab photonic crystal morphology emerged at any juncture in the phylogenetic tree, subsequent lineages appear to retain this trait of high order, closing a return to quasi-order or asymmetry (Fig. 1). This observation implies that the evolution of slab photonic crystal morphology imparts a selective advantage in a specific environmental context (Fig. 2). The exact nature of this advantage remains speculative at the current point.

To delve deeper into the potential biological functions of this observation, we turn our attention to the distribution and anticipated photonic properties associated with square and hexagonal lattice types. Although both lattice types are present in the Coscinodiscophyceae, the more recent Mediophyceae exhibit a pronounced constraint toward square lattice types. This observation suggests a potential correlation between lattice type and phylogenetic lineage. Upon scrutinizing the predicted photonic properties of the square lattice type, a particular pattern emerges. The anticipated spectral range of the photonic stopband predominantly aligns with the visible light spectrum in most cases (Fig. 3A). This alignment suggests a role in energy absorption in the visible spectrum of light, potentially for photosynthesis or processes related to sensing visible light. In contrast, hexagonal lattice types, characterized by larger lattice parameters, interact with the near and mid-infrared spectral range (Fig. 3B). Such properties may be associated with functions related to heat dissipation or the thermal regulation of cellular processes. While these hypotheses remain speculative at present, the intriguing correlation between lattice types, phylogeny, and distinct spectral ranges within the electromagnetic radiation spectrum prompts further exploration. The diverse spectral properties associated with different lattice types underscore that slab photonic crystals may bear different functionalities across diatom biodiversity, such as in light absorption and thermal regulation. We may conclude that such properties are derived frustule traits, as they emerge from quasi periodic templates.

The absence of photonic crystal candidates within the girdle elements of the Bacillariophyceae can be readily attributed to the ultrastructure inherent in the girdle elements. Many of the girdle elements in bacillariophycean taxa only bear a single row of pores or lack perforation entirely, particularly among the raphe-bearing pennates (Bacillariophycidae). A single pore line does not possess the necessary periodicity to create a photonic stopband, as a central property of the photonic crystal. The selective force, if any, driving this morphological shift is unknown. It might be tempting to relate the raphe (the slit on the valve face that serves as the basis of motility for this subclass) to this change, hypothesizing that the evolution of motility in the diatom cell meant the diatom could reorient itself to alter light availability by a secondary derived trait and no longer required photonic crystal capabilities in the girdle elements. However, this shift towards single pore rows and unperforated girdle elements already appears to begin in the araphid pennate diatoms (Fragilariophyceae), which lack raphid motility.

With regards to ecology, all the taxa with slab photonic crystal properties in the girdle elements are planktonic diatoms found in marine environments. In fact, *Trieres* is one of the few planktonic genera in the Odontellaceae family (though *Triceratium robsertsianum* is a benthic taxon). This dataset exhibits a notable bias towards marine benthic taxa, primarily during their greater availability during its construction. Additionally, it can be contended that there is also a strong bias in diatom diversity towards benthic taxa, given their presumed higher diversity and prevalence^24^.

It may be predicted that more of those slab photonic crystal structures are existent in yet unstudied species. Between any intraspecific variation in the genus *Coscinodiscus* and the slab photonic crystal morphology suggested in the additional genera presented here (*Proboscia* and *Trieres*), this could provide researchers a natural catalogue to select morphologies tailored to specific applications, precluding the cost and time required to engineer and fabricate similar nanomaterials. This study demonstrates the outstanding natural variation of slab photonic crystal morphologies across the diversity of diatoms, covering, with the resulting stopband properties, the full visible light to mid infrared spectral range of electromagnetic radiation. Why and under which environmental conditions slab photonic crystals evolved in the diatom frustule remain open questions; however, the presented data suggest that slab photonic crystal properties arose as derived frustule traits, as the presence of various lattice types (hexagonal or square) and the coexistence of quasi-photonic and well-defined slab photonic crystal morphologies are correlated with phylogenetic information. Importantly, the presented data exclude the possibility that natural slab photonic crystal morphologies emerged without functionality as random byproducts along the evolutionary trajectory of diatoms.

## Materials and Methods

### Morphological observations

The organic material was cleared from the frustules of existing diatom cultures at the University of Texas, Austin. Cells were treated in a 1:1:1 solution of cultured material:30% hydrogen peroxide:70% nitric acid for approximately 48 hours, centrifuged at 4500 rpm for 20 min (TX-400 rotor, Sorvall ST16R Centrifuge, Thermo Scientific). The pellets were then rinsed in distilled water and repeatedly centrifuged until neutral pH was achieved. The neutral solution was then pipetted onto 12 mm diameter cover glasses and dried overnight. Cover glasses were subsequently mounted onto aluminum stubs and coated in 15 nm of iridium or platinum/palladium using a Cressington 208 Bench Top Sputter coater. Scanning electron microscopy (SEM) was conducted using a Zeiss SUPR 40 VP scanning electron microscope (Carl Zeiss Microscopy, Thornwood, NH, USA).

### Identification of slab photonic crystal morphologies

The structural morphology of girdle bands was examined using size-calibrated SEM images. The SEM micrographs analyzed were from the four different taxonomic categories in an initial dataset containing Coscinodiscophyceae (n=44), Mediophyceae (n=132), Fragilariophyceae (n=95), and Bacillariophyceae (n=122). The corresponding categories were identified either with available DNA data or based on SEM morphological characterization.

The morphological analysis aimed to distinguish between square and hexagonal lattice types, and to eventually identify slab photonic crystal morphologies. The criteria included:

1. Exclusion of entries with girdle bands lacking pores or featuring only a single line of pores radiating in the x direction, or random distribution of porous features.
2. For entries with a periodic pore pattern in the x direction, determination of the number of pores in the z direction (see Fig. 4A). For identification, crystal morphologies required a periodic pore pattern in at least two dimensions, with exclusion criteria of fewer than 5 pores in the z direction. These were included in the identification of quasi-periodic morphologies. Specimens with more than 15 pores in each direction were classified as slab photonic crystals, considering the significance of pore number for photonic response.
3. The remaining SEM images from 2) underwent Fast Fourier Transform (FFT) image analysis using Fiji (ImageJ)^25^, a potent method converting spatial information into spatial frequency domains. FFT analysis provided quantitative information on the lattice parameters and allowed us to exclude those species showing a lack of symmetry on their FFT image. Lattice morphologies (square or hexagonal) were differentiated by analyzing the normalized modules of the two lattice vectors (a_1_ and a_2_) and the angle formed between them (φ). A square lattice is then defined by a_1_ = a_2_ and φ = 90^0^. Hexagonal lattices were defined as a_1_ equals a_2_, and the angle φ equals 120 degrees. In the case of a highly symmetric and orderly lattice of pores, multiple reciprocal lattice points exist within the FFT image. Each of these points corresponds to higher spatial frequencies, and their occurrence is more prominent as we move farther away from the central region of the image. The distance of each high intensity point to the center shows multiples of the periodicity of the structure in a given direction. In some cases, the lattice is very well-defined and the pores are well ordered, which reflects on several reciprocal lattice points over a uniform background. In certain scenarios, the FFT analysis reveals distinct symmetry points set against a pronounced circular symmetry background, particularly at low spatial frequencies. This observation suggests the presence of areas with short-range order in the image, albeit alongside elements that exhibit disorder. Consequently, we have categorized these cases as quasi-ordered square or hexagonal patterns (Fig. 1D). It is important to note that in this context, we have labeled as “quasi-ordered” only those instances where a clear pattern was discernible in the FFT analysis. It is possible that additional quasi-ordered patterns may exist within this dataset due to variations in image quality.

All entries that successfully passed this scrutiny underwent analysis for pore period (in the x direction) and pore diameter. Additionally, we classified the types of pores, differentiating between regular pores, elongated pores, sieve pores, or other ornamental structures (see Supplementary Dataset S1).

### Numerical analysis of photonic crystal properties

The structural parameters of candidates demonstrating FFT symmetry, as outlined above, were used to compute the central reflectance of the photonic stopband. This computation incorporated refractive index approximations derived from models established for slab photonic crystals in diatoms, as elaborated elsewhere^3^. It is important to acknowledge that the resulting central spectral reflectance values might not be entirely accurate, given the assumptions made during simulations, as observed in the case of species *Coscindodiscus granii*. However, considering that the spectral range is predominantly influenced by the parameter pore period, the anticipated realistic values are expected to align with these modeled data.

### Molecular phylogenetic analyses and ancestral state reconstruction

Among the strains used in the morphological analysis, nearly 300 had corresponding DNA sequence data from the nuclear-encoded ribosomal small subunit (nrSSU), and the chloroplast-encoded *rbc*L and *psb*C genes generated for molecular phylogenetic studies. The DNA extraction, PCR amplification and Sanger sequencing protocols for these data are covered in Lobban et al^26^. These loci were assembled into a concatenated dataset. All analyses for phylogenetic inference and ancestral state reconstruction were performed on the University of Alabama supercomputer cluster (UAHPC). Both Bayesian inference (BI) and maximum likelihood (ML) phylograms were inferred using ExaBayes^27^ and Iqtree^28^, respectively. For ancestral state reconstructions, data from comma separated value (csv) tables (Supplementary Dataset S2) were mapped onto the ML phylogram. For taxa with missing character state data in the csv file, the phylogram was pruned to match. Both discrete and continuous character sets were mapped via Phytools version 1.0-3 package^29^ as implemented through R version 4.2.0^30^. Model optimization for discrete character states were analyzed using the *fitMk* function in Phytools, that fits the *Mk* model^31^ to a phylogenetic tree and a discrete character state matrix under the equal rates (ER), symmetric rates (SYM), or all rates different (ARD) reversible models. The best model was selected by a comparison of Akaike information criterion (AIC) scores (Supplementary Table S1). The *make*.*simmap* function (Phytools) was used in *parallel* (R) to stochastic map character states with a resulting total of 100,000 simulations for each csv file (http://blog.phytools.org/2017/11/running-makesimmap-in-parallel.html). For binary discrete character states, density maps were constructed using the *densityMap* function in Phytools.

## Acknowledgments

JWG is grateful for support through a Concurso Estímulo ao Emprego Científico Individual grant (no. 2020.04217.CEECIND) from the Fundação para a Ciência e a Tecnologia (FCT), Portugal. JWG and MLG thank the FCT for their support through grant no. PTDCBTA-BTA20612021 (NASCADIA project). The work by MLG was also supported by the ENIGMA project (grant code 342255) funded by the Research Council of Norway. JWG also thanks the FCT/MCTES for the financial support to CESAM (UIDB/50017/2020+UIDP/50017/2020+LA/P/0094/2020). MPA is grateful to the following researchers for their assistance in collecting material and cultures for electron microscopy and sequencing: E. Theriot, D. Williams, A. Witkowski, J. Witkowski, C. Lobban, R. Majewska, T. Frankovich, P. Sims, P. Kociolek, R. Jordan, S. Sato, A. Alverson, M. Sullivan, S. Bosak, J. S. M. Sabir. DWL thanks Liam J. Revell for his assistance in mapping character trait data to the phylogeny and Michael R. McKain for his insightful discussions on the same topic. D.W.L. is grateful to the University of Alabama Supercomputing Cluster (UAHPC) for computational time used for phylogenetic inference and character trait mapping.

